# Optimization of capture protocols across species targeting up to 32000 genes and their extension to pooled DNA

**DOI:** 10.1101/2022.01.10.474775

**Authors:** Cédric Mariac, Kévin Bethune, Sinara Oliveira de Aquino, Mohamed Abdelrahman, Adeline Barnaud, Claire Billot, Leila Zekraoui, Marie Couderc, Ndjido Kané, Alan Carvalho Andrade, Pierre Marraccini, Catherine Kiwuka, Laurence Albar, François Sabot, Valérie Poncet, Thomas LP Couvreur, Cécile Berthouly-Salazar, Yves Vigouroux

**Affiliations:** DIADE, Univ Montpellier, IRD, CIRAD, Montpellier, France; Federal Univ of Lavras, Brazil; Rice Research and Training Center, Field Crops Research Institute, Agricultural Research Center, Kafrelsheikh, Egypt; AGAP Institut, Univ Montpellier, CIRAD, INRAE, InstitutAgro-SupAgro, Montpellier, France; CIRAD, UMR AGAP Institut, F-34398 Montpellier, France; LMI LAPSE, ISRA, IRD, Dakar, Senegal; EMBRAPA Coffee-INOVACAFE, Lavras, Brazil; CIRAD – UMR DIADE, F-34398 Montpellier, France; NARO, Kampala, Uganda; Wageningen University and Research Centre, Wageningen, Netherlands; PHIM Plant Health Institute of Montpellier, Univ Montpellier, IRD, CIRAD, INRAE, Institut Agro, Montpellier, France

**Author notes:** Yves Vigouroux Institut de Recherche pour le Développement, Univ Montpellier, Unité Mixte de Recherche Diversité Adaptation et Développement des Plantes (UMR DIADE), Montpellier, France.

## Abstract

**Premise:** In-solution based capture is becoming a method of choice for sequencing targeted sequence.

**Methods and results:** We assessed and optimized a capture protocol in 20 different species from 6 different plant genus using kits from 20,000 to 200,000 baits targeting from 300 to 32,000 genes. We evaluated both the effectiveness of the capture protocol and the fold enrichment in targeted sequences. We proposed a protocol with multiplexing up to 96 samples in a single hybridization and showed it was an efficient and cost-effective strategy. We also extended the use of capture to pools of 100 samples and proved the efficiency of the method to assess allele frequency. Using a set of various organisms with different genome sizes, we demonstrated a correlation between the percentage of on-target reads *vs*. the relative size of the targeted sequences.

**Conclusion:** Altogether, we proposed methods, strategies, cost-efficient protocols and statistics to better evaluate and more effectively use hybridization capture.

## INTRODUCTION

The reduced representation library approach (RRB) has become a widely used tool in molecular ecology and phylogeny. Some approaches are based on DNA restriction, e.g. RAd-seq (Andrews et al., 2016) or genotyping-by-sequencing (Elshire et al., 2011; GBS), in which the studied polymorphism is not targeted. In other approaches hybridization capture is used notably in phylogenetic studies (Mandel et al., 2014, Weitemier et al., 2014, Kollias et al., 2015, Nicholls et al., 2015, Stephens et al., 2015a, Stephens et al., 2015b, McCartney-Melstad et al., 2016, Portik et al., 2016, Couvreur et al., 2019) and to a lesser extend to diversity studies (Asan et al., 2011, Rosani et al., 2014, Kistler et al., 2015). Among these approaches, in-solution hybridization with DNA or RNA probes are the most frequent. Although they are common in human genetic studies (Asan et al., 2011), there are still underused in molecular ecology.

To assess the success of in-solution hybridization, the main parameter is “fold increase”, *i*.*e*. to what extent a target sequence is enriched compared to the rest of the genome. This statistic is referred to here as x-fold enrichment. It measures how effective the enrichment is in terms of increasing the proportion of the target. Another parameter is the percentage of reads mapping on the target sequence (on-target reads, in contrast to off-target reads). This parameter is useful to assess the cost effectiveness of the experiment. Both statistics, x-fold enrichment and the percentage of on-target reads, are used interchangeably in the literature, whereas in fact, they assess two different aspects of the capture protocol. Such an enrichment protocol allows the analysis of many individuals, which is extremely useful in population genetics or phylogenetic studies. In addition, the combining of several individuals in a single hybridization may further reduce the cost. Lastly, pooled DNA sequencing is also increasingly used to assess allele frequency in a population (Pool-seq, Futschik and Schlötterer, 2010). In this approach, several individuals of the same population are sequenced together to estimate population allele frequency (Futschik and Schlötterer, 2010). The usefulness of the pool-seq approach combined with capture have not yet been fully evaluated yet, but is a promising avenue for future study.

Here, we used a set of 21 plant species with different genome sizes and target sequences to optimize in-solution capture approaches. X-fold enrichment and the percentage of on-target reads were calculated to assess whether the approach is really both cost and experimentally effective. We show that high multiplexing for a single hybridization is both efficient and cost effective. A further extension of this approach was developed for pooled DNA sequencing. Finally, we demonstrated a relationship between the size of the target sequence relative to the size of the genome and the percentage of on-target reads, allowing better design and efficient prediction in any specific experiment.

## MATERIALS AND METHODS

### Plants materials

A total of 21 species from four families belonging to 13 genera (Annonaceae: *Anaxagorea, Annickia, Anonidium, Greenwayodendron, Monanthotaxis, Monocarpia, Monodora, Neouvaria;* Arecaceae: *Podococcus;* Poaceae: *Cenchrus, Digitaria, Oryza;* Rubiaceae: *Coffea)* were used in this study (Table S1a). Fresh leaves, dried leaves and tissues collected in herbarium were used for total DNA extraction following a previously described protocol (Mariac et al., 2006). DNA was extracted from individual plants. In addition, for pearl millet variety PE5487, leaves from 100 plants were bulked before DNA extraction using a poolseq approach.

### Locus target selection and bait design

Seven enrichment kits were tested (Table S1b). Five of the seven kits were specific to the present study and only two have already been described (*Arecaceae*: Heyduck et al., 2016; *Annonaceae*: Couvreur et al., 2019). The total size of the targets to be captured varied from 204 kb to 12 Mb. RNA baits designed varied from 80bp to 120bp and from a 3X to 0.5X tiling (Table S1b). The targets were exonic sequences available either from transcriptome assemblies for *Cenchrus* and *Digitaria* (Sarah et al., 2016), or in the case of *Cenchrus americanum* (Varshney et al., 2017), *Oryza sativa* (Kawahara et al., 2013), and *Coffea canephora* (Denoeud et al., 2014), from fully annotated genomes. Biotinylated baits were designed and synthesized by Mycroarray (Ann Arbor, Michigan, USA). Mycroarray ran the set of baits through RepeatMasker v4,0 (Smit et al. 2013) over the closest available reference genome (Table S1b) to avoid designing baits that target repetitive sequences.

### Library preparation and sequencing

Libraries were prepared according to the protocol of Mariac et al. (2018). Briefly, DNA samples were sheared to yield 400-bp fragments. DNA was then repaired and tagged using 6-bp barcodes to allow further multiplexing. Real-time PCR was performed to generate ready-to-load libraries. These libraries were either immediately sequenced for a shotgun genomic sequencing or enriched by capture according to the Myselect protocol (Mycroarray) before sequencing.

We tested different multiplexing levels for maximum possible cost reduction. From 1 to 96 equimolar libraries were multiplexed in a single DNA capture reaction. We then assessed the impact of multiplexing on enrichment efficiency.

Using the poolseq protocol, we built 100 individual libraries from 100 individual plants, and two libraries made by bulking exactly the same 100 plants. One (mock) library corresponded to an equimolar mix of the DNAs extracted from the 100 individuals, while the other library corresponded to a single extraction of DNA from the pooled leaves of the same 100 individuals.

A total input of 500 ng DNA was used per capture and hybridization was performed at 65 °C for 18 h including blocking oligonucleotides with six inosines at the barcode location to reduce unspecific hybridization. The immobilization and washing steps were conducted as recommended by the supplier. After probe-target hybridization (at 98 °C for 5 mins) the resulting enriched libraries were amplified using the KAPA Biosystem Real Time PCR Kit (KK8221) according to the supplier’s recommendations. Paired-end sequencing was performed on an Illumina MiSeq (2×150 bp at CIRAD, Montpellier, France or on the Hiseq2000 platform (Genotoul, Toulouse).

### Bioinformatics analysis

demultiplexing, data cleaning, mapping, SNP calling. Demultiplexing based on 6-bp barcodes was performed using a Python script DEMULTADAPT (https://github.com/Maillol/demultadapt), using a 0-mismatch threshold. Adapters and low-quality bases were removed using CUTADAPT 1.8 (Martin, 2011) with the following options: quality cut-off = 20, minimum over-lap = 7 and minimum length = 35 (Table S1c). Reads with a mean quality lower than 30 were discarded using a PERL script (https://github.com/SouthGreenPlatform/arcad-hts/blob/master/scripts/arcad_hts_2_Filter_Fastq_On_Mean_Quality.pl). Mapping was performed using bwa mem 0.7.5a-r405 (Li et al., 2009) and the selected targets as the reference. GATK v3.3-0-g37228af UnifiedGenotyper (McKenna et al. 2010) was used for SNP calling. For the poolseq approach, we only kept SNPs with no missing data with a minimum coverage of 100 across the 100 individuals and at least 50 reads per bulk. Allele frequencies were extracted from the VCF files using VCFtools v 0.1.14 (Danecek et al., 2011). For the poolseq bulk, allele frequencies were calculated based on the count of the reference and alternate alleles using VCFtools v 0.1.14 (Danecek et al., 2011).

### Estimation of enrichment efficiency

We first calculated the percentage of on-target reads, *i*.*e*. the number of reads mapped on the target references divided by the total number of reads. We then calculated the x-fold enrichment (x-fold), *i*.*e*. the number of on-target reads after enrichment divided by the number of on-target reads without enrichment. The latter number was estimated based on sequencing of the whole genome (Table S1d). These two statistics were calculated for each library; averages and standard deviations were calculated per species and per multiplex (Table S1d).

### Literature review

To gain comprehensive insight into the methodology, we compared our results with those obtained in previously published studies. Articles that described the use of a similar method on fresh and ancient DNA of plants and animals were downloaded and analyzed (see list of publications in supplementary file). For each study, when possible, we re-encoded or calculated both statistics, *i*.*e*. the x-fold enrichment and the percentage of on-target reads. For each of these studies, we also calculated the ratio of the total size of the targeted sequences to the total genome size. When necessary, the genome size was evaluated based on the litterature C-value (Dolezel et al., 2003, detailed in table S2).

## RESULTS AND DISCUSSION

### Results of multiplexing did not change either the x-fold or the number of on-target reads

We first analyzed the results obtained on three species in greater detail (Table S1d): *Cenchrus americanus*/*Pennisetum glaucum* (kit MIL-328), *Digitaria exilis* (Kit Fonio) and *Coffea canephora (*kit Coffee).

For pearl millet, with a capture approach targeting 328 genes (MIL-328), the percentages of on-target reads after enrichment were 14.3 % (se=3.34), 17.3 % (se=0.21) and 18.4 % (se=0.16) for 1, 8 and 30 libraries per capture, respectively (Figure S1, Table S1d). A slightly higher number of on-target reads was retrieved with higher multiplexing (Kruskal-Wallis, H=12.38, *p*=0.002). The x-fold enrichment associated with multiplexing varied from 78.9, 99.3 and 106.6 for 1, 8 and 30 libraries per capture, respectively (Figure S1). Enrichment was more efficient with higher multiplexing (Kruskal-Wallis, H=17.82, *p*<0.0001).

For *Digitaria exilis*, with a capture targeting 3000 genes, the average percentages of on-target reads after enrichment were 83.5 % (se=0.009), 83.7 % (se=:0.851) and 85.3 % (se=1.55) for 1, 8 and 39 libraries per capture, respectively. The percentage of useful reads was slightly better with higher multiplexing (Kruskal-Wallis, H=8.19, *p*=0.016). The average x-fold values were similar (Kruskal-Wallis, H=4.01, *p*=0.134) with 33.6, 35.7 and 34.5 for 1, 8 and 30 libraries per capture, respectively (Figure S1).

For *Coffea canephora*, the targeted 323 genes (de Aquino et al. 2021) represented a total length of 1.3 Mb *i*.*e*. 0.2% of the whole genome (1C = 710 Mb). The percentage of reads on-target after enrichment of 70%. The x-fold level of target enrichment was 270 compared to a non-enriched genome sequencing, and only slightly lower than a single captured library x-fold enrichment (320).

Our results showed that the percentage of on-target reads was generally not affected by the number of individuals multiplexed for hybridization across the other kits and species (Figure S2). The percentage ranged from 15% for a single capture on pearl millet (MIL-358) to 80% on rice (Figure 1, Figure S2). The x-fold enrichment ranged from 5 on pearl millet to 400 for *Annonaceae* (Figure S2). We tested up to 96 individuals captured at once (Figure 1, Figure S2). Based on an estimated cost of USD3,600 for 16 captures (https://arborbiosci.com/products/custom-target-capture/), the capture price per sample amounted to USD225 without multiplexing, and to only to USD2,34 with multiplexing of 96 samples, as proposed in our protocol, which makes the cost really affordable. Our result thus showed that multiplexing several individuals in a single capture hybridization is an efficient and cost-effective strategy.

**Figure 1.**
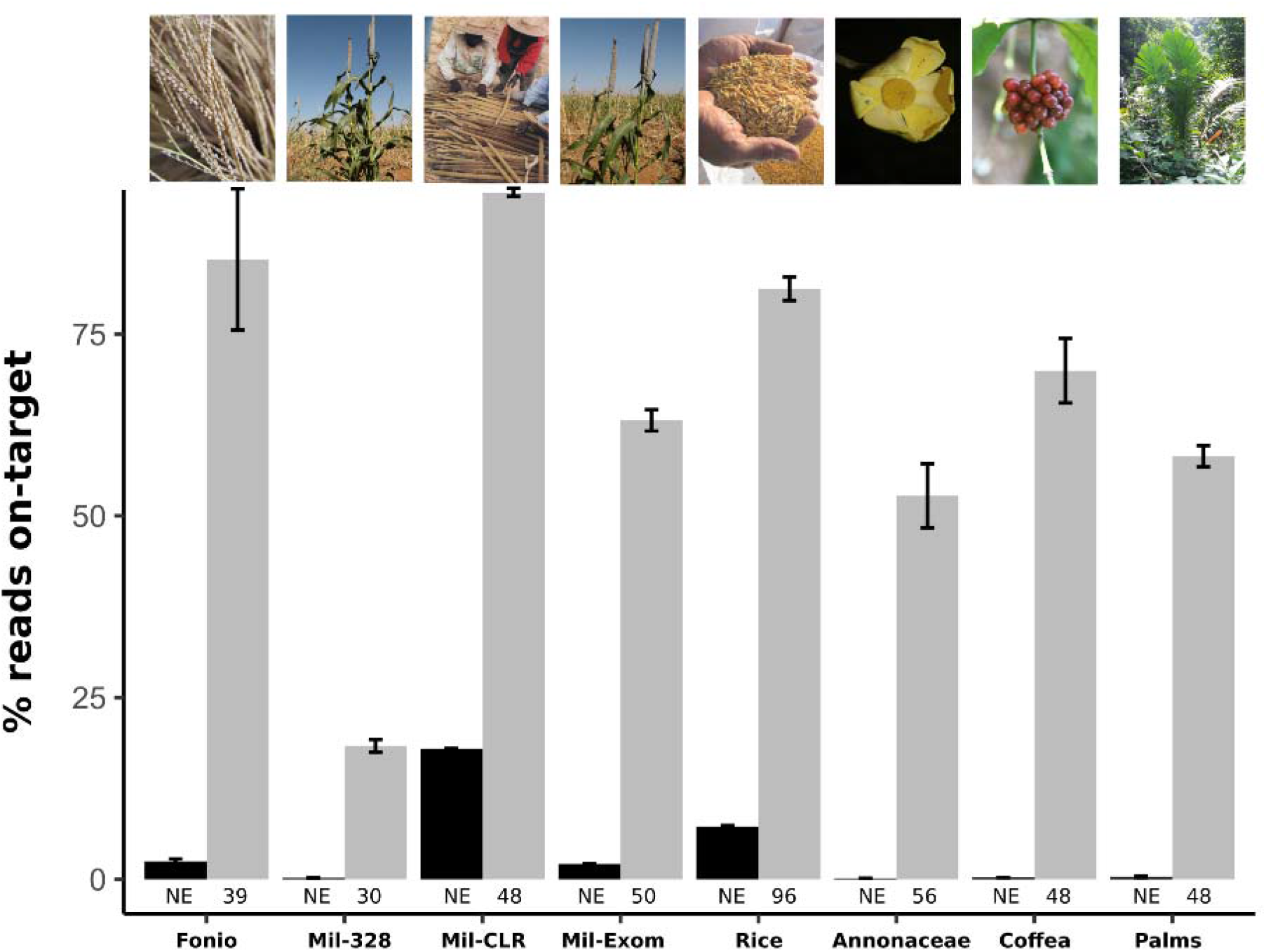
Percentage of on-target reads across species and capture design. Percentage of on-target reads (grey bars) compared to unenriched libraries (dark bars) and their confidence intervals. For each. the species studied are *Digitaria exilis* (fonio), *Cenchrus americanun* (with three different kit: MIL-328, MIL-CLR, MIL-EXOME, see Table SXX), *Oryza* spp. (Rice), *Annonacea* spp. (Annonaceae), *Coffea* spp. (Coffea), several palm species (Palms).

### Analysis of the poolseq approach with capture

To evaluate the accuracy of the capture approach for poolseq DNA, we compared the allele frequencies obtained from libraries performed on 100 individual DNA samples, on a bulk of 100 DNA samples (bulk 1), and on a DNA sample extracted from the pooled leaves of 100 individuals (bulk 2). We identified a total of 62,320 SNPs in the 100 individuals and two bulk samples. The allele frequencies between these two bulk samples (r^2^=0.99, *p*<10^−20^) were highly correlated, as well as between individual libraries and the bulks (r^2^=0.98, *p*<10^−20^ between individual libraries and bulk 2) (Figure 2). Capture thus made it possible to effectively retrieve the allele frequency of a bulk of individuals. Our protocol makes this approach more broadly accessible, even for large genomes. Such an approach could also be used for bulk segregant analysis (Takagi et al., 2013) to identify functional variations linked to specific traits.

**Figure 2.**
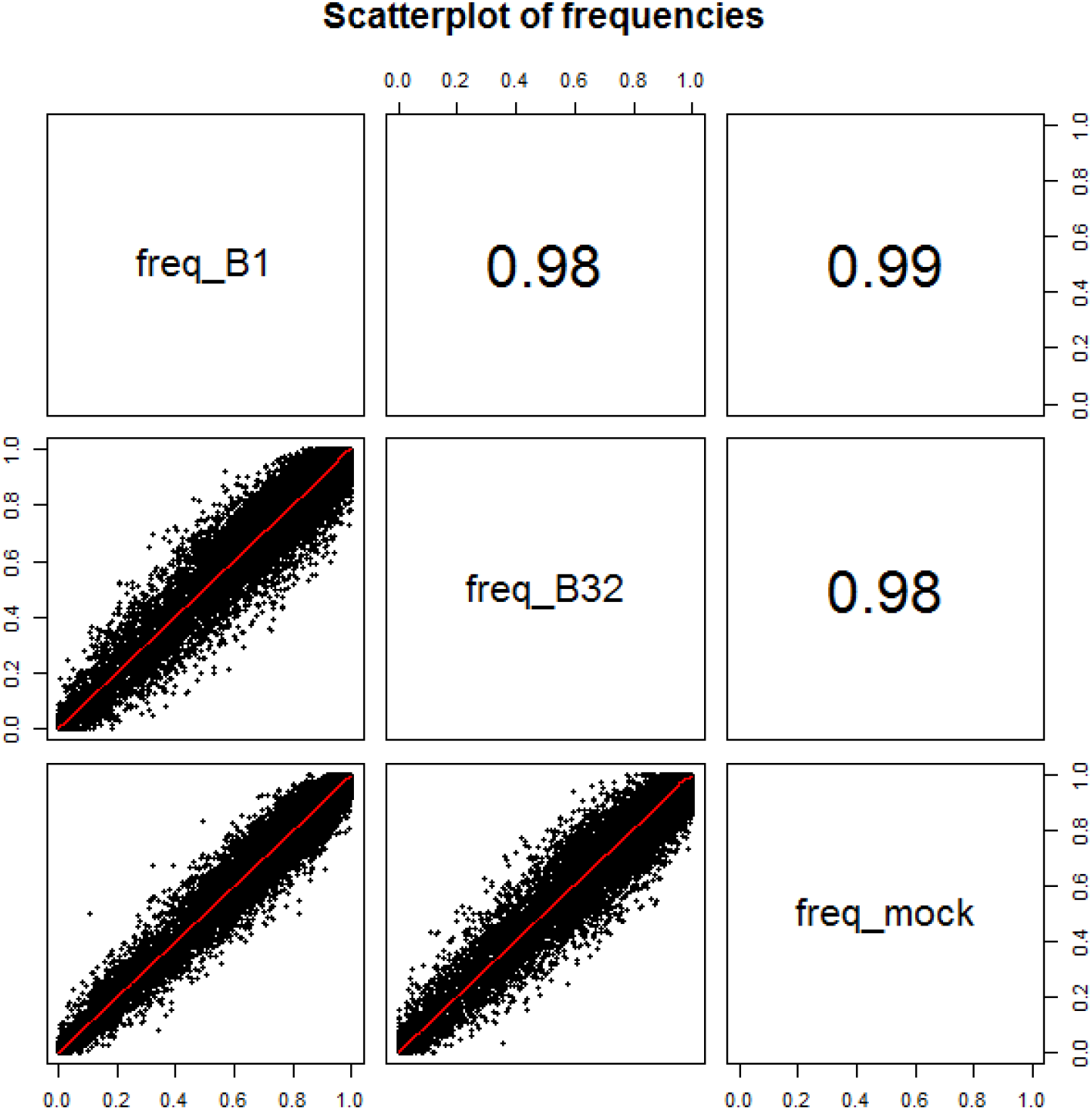
Correlation of expected and observed allele frequencies using poolseq capture protocols. Calculated correlations between allele frequencies estimated based on individual capture of 100 individuals (Freq_B1), and on capture of bulk samples. One bulk sample was made of an equimolar concentration of DNA of 100 individuals (freq_mock), the other was made of a DNA extract of 100 pooled pieces of leaves from the same 100 individuals (freq_B32). The correlations were highly significant between all the three experimental conditions (r^2^≥0.98, *p*<10^−20^).

### Literature review and meta-analyses

We retrieved or calculated the percentage of on-target reads from 22 studies (Table S2). Using these data and the seven studies presented here, we evidenced a logarithmic relationship (r^2^=0.54, *p*<2.10^−5^) between the percentage of on-target reads and the relative size of the targeted sequence (Figure 3). For small targets representing less than 1% of the genome, it is not uncommon to find a relatively low percentage of on-target reads (Figure 3), but the percentage of on-target reads increases exponentially up to almost 100% with an increase in the relative size of the target. This relationship makes it possible to predict the percentage of on-target reads based on the size of the genome and the size of the targeted sequence, meaning a preliminary assessment of the cost effectiveness of the experiment can be performed before design. An extreme example of capture design that targets a very short sequence (∼500bp) will lead to very low on-target reads percentage, with 1% or less (Maggia et al., 2017). Recent results show that the protocol and two rounds of capture can be performed to increase on-target reads on target up to 70% (Mariac et al., 2018). With the low cost of capture for high multiplexing, two rounds of capture would be very cost effective.

**Figure 3.**
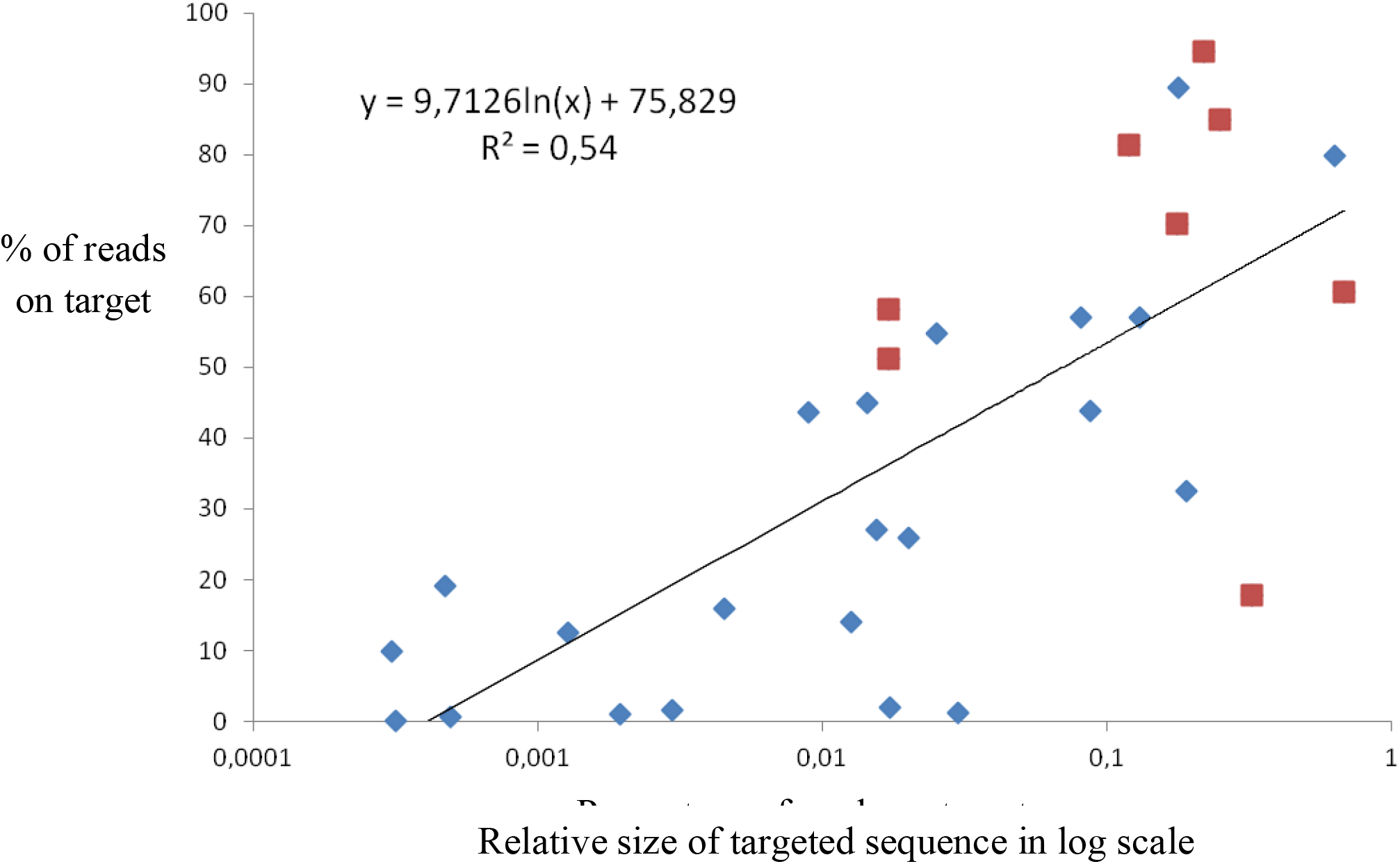
Relationship between the percentage of on-target reads and the relative size of the targeted sequence. Significant relationship between the logarithm of the relative targeted sequence (size of the target divided by the size of the genome) and the number of on-target reads (r^2^=0.54, *p*<2 10^−5^). Red dots represent data collected in the present study and blue dots represent data retrieved from the literature (Table S2).

Only 5 out of 22 (23%) studies in the literature provided the x-fold enrichment level. It was thus rather difficult to perform meta-analysis on this statistic. X-fold enrichment really measures the effectiveness of the experiment because it assesses how many more sequences are obtained after capture compared to a non-enrichment baseline. We suggest that both the percentage of on-target reads and x-fold enrichment should be more frequently reported in the literature. When the x-fold enrichment is not reported, it is difficult to really assess if and how well the capture experiment worked.

## CONCLUSION

We assessed capture protocols in different plant species, and showed high multiplexing per capture is an efficient strategy across species. We demonstrated that this capture strategy can easily be extended to pooled DNA samples to assess allele frequency. We highlighted a relationship that allowed the percentage of on-target reads to be estimated. The early estimation of this important statistic could guide the protocol and the size of the target sequence for more efficient use of these approaches. Finally, we highlighted the importance of reporting both the percentage of on-target reads and the x-fold enrichment in any capture experiment to allow better assessment of the approach.

## ACKNOWLEDGEMENT

This work was funded by an ANR grant to YV; CBS, CM, YV, FS and by Agropolis Foundation (Reference ID 1402-003) through the « Investissements d’avenir » program (Labex Agro:ANR-10-LABX-0001-01). VP, SOA, CK, AA and PM were supported by two Agropolis Foundation - CAPES (Coordenação de Aperfeiçoamento de Pessoal de Nível Superior) projects, i.e. ID 1002-009 and ID 1402-003 (CLIMCOFFEA), through the Investissements d’avenir program (Labex Agro:ANR-10-LABX-0001-01), in the framework of I-SITE MUSE (ANR-16-IDEX-0006). MA had a fellowship from the French Embassy in Egypt, Institut Français d’Egypte and the Science and Technology Development Fund. The authors acknowledge the ISO 9001 certified IRD itrop HPC (member of the South Green Platform) at IRD Montpellier for providing HPC resources that have contributed to the research results reported within this paper. URLs: https://bioinfo.ird.fr/ and http://www.southgreen.fr. We thanks Anaïs Dequincey and Coralie Picard for help in developing some of the experiments.

## Data availability

Sequencing reads were deposited in the National Center for Biotechnology Information Sequence Read Archive (BioProject ID: PRJNA431698, BioSample accessions: SAMN08625127-SAMN08625640).

## Supporting information

Fasta files of baits and target used for each kit:

TARGET-MIL328.fasta

TARGET-MIL-CLR.fasta

TARGET-palms.exons.final.fasta

TARGET-RICE.fasta

bait-Annonaceae_nuc.fas

baits- MIL-EXOME-bait-80-160-first7-fixed.fas

baits-COFFEE.fas

baits-MIL-328.fas

baits-MIL-CLR.fas

baits-PALM_EXONS.fas

baits-RICE.fas

TARGET-MIL-EXOME-152169_stringent_baits-coordv1.1.bed

TARGET-Annonaceae_nuc.fasta

TARGET-COFFEE.fasta

TARGET-FONIO.fasta

## Supplementary figures

**Figure S1.**
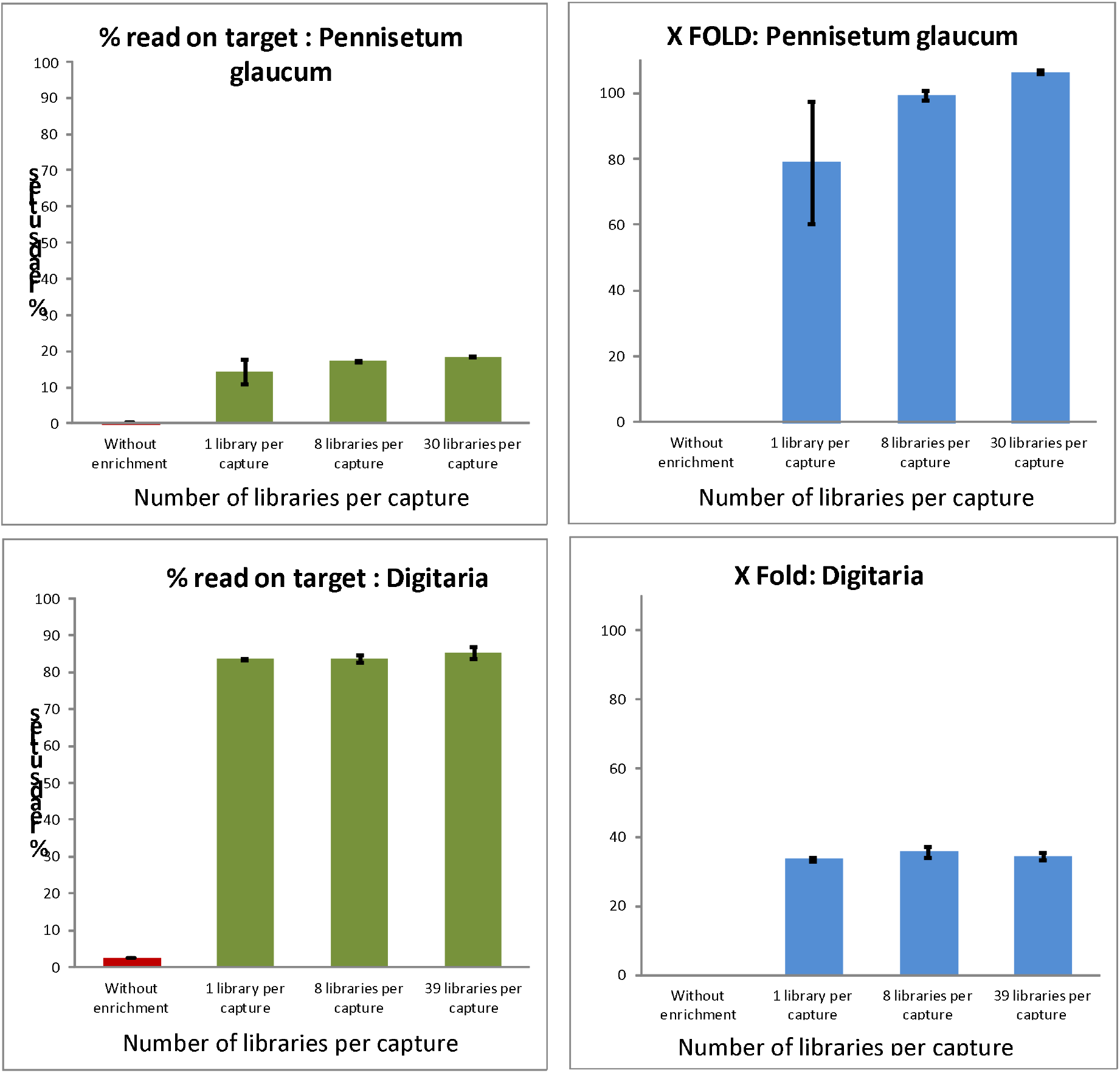
X-fold enrichment and percentage of on-target reads for *Pennisetum glaucum* and *Digitaria exilis*. We tested different multiplexing of samples for a single capture experiment (1, 8 and 39 libraries per capture for *Pennisetum glaucum*; 1, 8 and 39 libraries per capture for *Digitaria exilis*). Experiments included non-enriched libraries as controls. Both the fold enrichments and the percentages of on-target reads obtained with different multiplexing were very similar and even tended to be slightly better with an increase in the number of samples.

**Figure S2.**
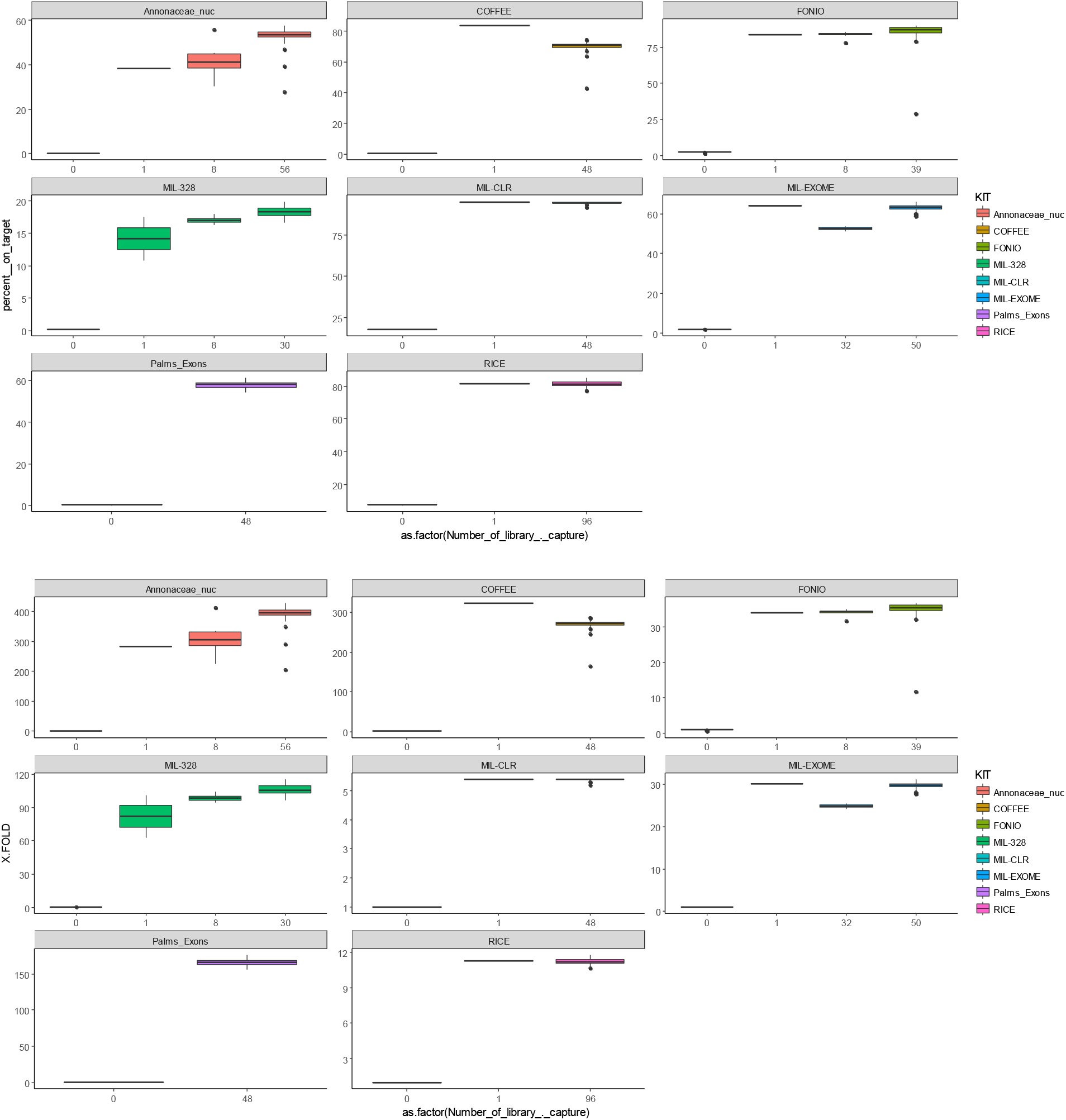
X-fold and percentage of on-target reads for all the species. For all the species (7 different capture kits), we tested different multiplexing of samples for a single capture experiment. The experiment included non-enriched DNA sequence as a control.

**Table S1. Passport data, bait design, bioinformatics and raw mapping data**.

*Table S1a. passport data of all samples used in this study*.

The table list : plant name, the collection ID of the accession, the species name, type, origin of the material and the country where the accession was sampled.

*TableS1b. Details of the kit used in this study*.

*Table S1c. Option used for bioinformatic analysis and mapping reads*

*Table S1d. List of individual and each sequencing files*

For each individual, its accession name (Accession_IR, Accession_ID2), the species, the tag used for sequencing (internal information RUN-index-TAG), an experimental code (Code), the library preparation (capture or direct shotgun without capture) and the kit used for the experiment are given.

We also give the number of libraries used for a single capture experiment, the number of Illumina reads, the number of reads mapping on target, the percentage of read on target, the x-fold enrichment

We also give reference of each sample in GenBank (Biosample identification, link and name of the fastq files)

see excel files

**Table S2.**
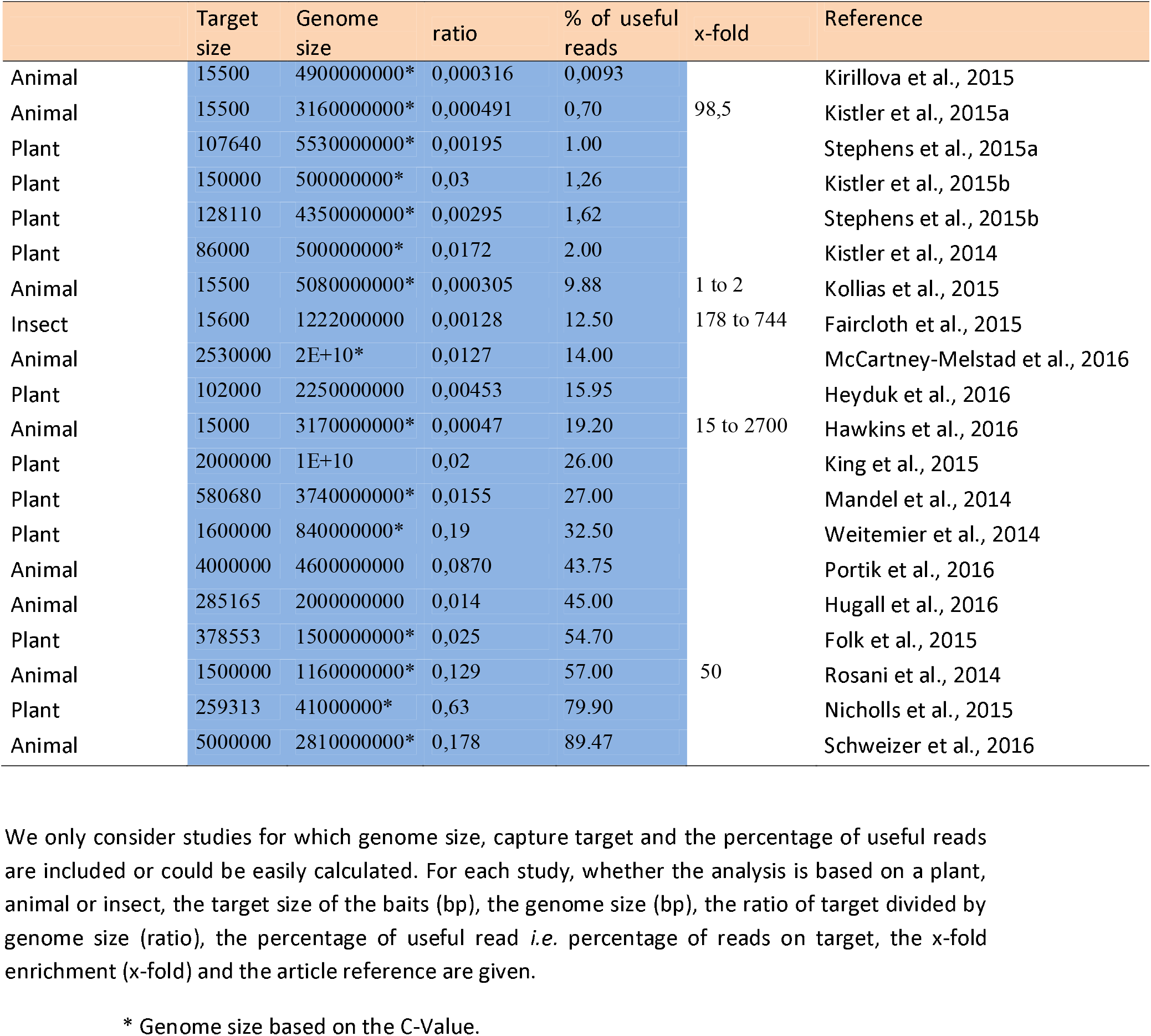
List of studies in which genome size, capture target and the percentage of useful reads are included.

